# Image-Informed Inverse Finite Element Analysis Reveals Altered Constitutive Behavior Following Controlled Uterine Tissue Remodeling

**DOI:** 10.64898/2026.08.23.746519

**Authors:** Mahmuda Raakib Arshee, Callan M. Luetkemeyer, Indrani C. Bagchi, Ayelet Ziv-Gal, Jodi A. Flaws, Adira Safar, A. J. Wagoner Johnson

**Affiliations:** Mechanical Science and Engineering, Grainger College of Engineering, University of Illinois at Urbana-Champaign, Urbana, IL, 61801, USA; Comparative Biosciences, College of Veterinary Medicine, University of Illinois at Urbana-Champaign, Urbana, IL, 61802, USA; Biomedical and Translational Sciences, Carle Illinois College of Medicine, University of Illinois at Urbana-Champaign, Champaign, IL 61820, USA; Carl R. Woese Institute for Genomic Biology, University of Illinois at Urbana-Champaign, Urbana, IL, 61801, USA; Biohub Chicago, LLC, Chicago, IL, 60642, USA; Beckman Institute for Advanced Science and Technology, University of Illinois at Urbana-Champaign, Urbana, IL, 61801, USA; Department of Bioengineering, Grainger College of Engineering, University of Illinois at Urbana-Champaign, Urbana, IL, 61801, USA; Materials Research Laboratory, University of Illinois at Urbana-Champaign, Urbana, IL, 61801, USA

**Keywords:** Uterus, glutaraldehyde crosslinking, inflation testing, inverse finite element analysis, Gasser-Ogden-Holzapfel model, micro-computed tomography

## Abstract

**Purpose:** Fibrotic remodeling of the uterus, associated with aging, disease, and environmental exposures, alters collagen organization and tissue stiffness, yet how these changes influence organ-level mechanical behavior remains poorly understood. Glutaraldehyde (GA)-induced collagen crosslinking was used as a controlled surrogate for fibrotic remodeling to determine whether image-informed inverse finite element analysis (iFEA), combined with inflation testing and micro-computed tomography (microCT), could detect and quantify the resulting changes in uterine constitutive behavior.

**Methods:** Murine uteri (n = 6 untreated, n = 6 GA-crosslinked) underwent volume-controlled balloon inflation with simultaneous microCT imaging to quantify deformation of the inner and outer wall boundaries for iFEA. Specimen-specific Gasser-Ogden-Holzapfel (GOH) finite element models were optimized by adjusting model parameters to reproduce experimentally measured wall contours throughout inflation. Model performance was evaluated using contour root mean square error (RMSE), and parameter identifiability was assessed through sensitivity analyses.

**Results:** GA treatment significantly increased inflation work, linear stiffness, and maximum inflation resistance (p < 0.001). The iFEA framework accurately reproduced experimental deformation (RMSE < 3%) and revealed significant increases in the estimated GOH parameters *C*_10_ (9.2-fold), *k*_1_ (2.0-fold), and *k*_2_ (2.7-fold), consistent with increased effective tissue stiffness and a shift toward earlier collagen fiber recruitment. Sensitivity analyses demonstrated unique, well-defined minima for all parameter combinations.

**Conclusion:** Image-informed iFEA provides a quantitative framework for relating collagen remodeling to organ-level uterine mechanics through specimen-specific constitutive parameter estimation. This approach establishes a foundation for investigating the mechanical consequences of uterine fibrosis and other remodeling processes.

## 1. Introduction

Collagen-rich soft tissues exhibit nonlinear mechanical behavior due to their hierarchical structure and the progressive recruitment of collagen fibers under load [1, 2]. Their characteristic response includes a compliant toe region followed by strain stiffening, which plays a critical role in the physiological function of soft organs, including the uterus [3, 4]. The uterine wall has distinct layers that are fiber-reinforced and in which the mechanically dominant myometrium comprises smooth muscle cells embedded within an extracellular matrix rich in collagen, elastin, and proteoglycans, with collagen fibers preferentially aligned in the circumferential and longitudinal directions [5–8]. Alterations to this collagen network, such as those induced by aging, disease, or environmental toxicant exposure, as examples, can significantly alter tissue mechanics and, consequently, organ function [9–16].

Despite the recognized importance of collagen organization in governing tissue mechanics, the relationship between uterine microstructure and organ-level mechanical behavior remains poorly understood. Although structural remodeling can be identified using imaging and histological approaches, these techniques do not quantify the mechanical consequences of such changes [17–22]. As a result, quantifying the mechanical consequences of structural remodeling remains a significant challenge.

Mechanical characterization of the uterus presents unique challenges because physiological loading occurs through pressure-driven distension of the uterine lumen, a loading mode that differs fundamentally from those employed in conventional mechanical testing [23, 24]. Accordingly, inflation-based protocols, also referred to as distension-based protocols, have been developed to preserve native tissue geometry while reproducing physiologically relevant multiaxial loading [25, 26]. These inflation-based methods have been successfully applied to characterize organ-level mechanical behavior and remodeling-associated changes in the arterial wall [27], murine vagina [25], cervix and uterus [28], lower urinary tract [29, 30], and GI tract [31, 32].

While inflation experiments characterize organ-level mechanical behavior, they do not directly identify the constitutive parameters governing the observed response. More recently, constitutive modeling coupled with inverse finite element analysis (iFEA) has been used to estimate material properties of human uterine tissue from indentation and uniaxial tension experiments [33–35]. However, these approaches have not yet been applied to pressure-driven inflation of uterine tissue. Integrating inflation experiments with iFEA therefore provides a means to relate organ-level mechanical behavior to the constitutive parameters governing uterine inflation.

A constitutive framework is required to relate experimentally measured deformation to the underlying constitutive behavior of the tissue. Anisotropic hyperelastic constitutive formulations such as the Gasser–Ogden–Holzapfel (GOH) model describe tissue behavior using parameters associated with matrix stiffness, fiber orientation, fiber stiffness, and fiber recruitment [36, 37]. When combined with experimentally measured deformation and specimen-specific geometry, iFEA can estimate constitutive parameters that provide insight into how tissue constituents govern mechanical behavior [38], as demonstrated across a range of soft tissues including the lung [39], sclera [40], pelvic floor muscle [41], arterial wall [27], myocardium [42], and skin [43].

To enable constitutive parameter estimation from uterine inflation experiments, we developed an image-informed iFEA framework that integrates balloon catheter–based inflation testing, micro–computed tomography (microCT) imaging, and specimen-specific finite element modeling. Experimentally measured tissue deformations acquired during controlled inflation were used to estimate GOH constitutive parameters, enabling specimen-specific characterization of uterine mechanical behavior. This framework provides an opportunity to determine whether controlled microstructural remodeling produces measurable changes in constitutive parameter estimates.

To evaluate the ability of this framework to detect microstructural remodeling, we employed glutaraldehyde-induced crosslinking as a controlled perturbation of the uterine collagen network. Glutaraldehyde (GA) is widely used as a collagen crosslinking agent and has been shown to alter the mechanical behavior of collagen-rich tissues in a predictable manner [44–46]. Although distinct from biological fibrosis, GA induces collagen crosslinking and increases tissue stiffness. Similar increases in tissue stiffness may arise following fibrotic remodeling associated with environmental toxicant exposure [9, 10]. Consequently, GA treatment provides a controlled means of introducing a large, reproducible alteration in tissue stiffness and serves as an initial validation of the ability of the iFEA framework to detect mechanically relevant changes in tissue behavior.

The objectives of this study were to (i) quantify uterine deformation during controlled inflation using micro-CT imaging, (ii) estimate specimen-specific constitutive parameters using image-informed iFEA, and (iii) determine whether the estimated parameters reflect mechanical changes indicative of collagen crosslinking. We hypothesized that GA-induced collagen crosslinking would alter the mechanical response of the uterus, resulting in systematic changes in the estimated GOH constitutive parameters. We further hypothesized that the image-informed iFEA framework would detect these changes, thereby demonstrating its ability to identify constitutive alterations associated with remodeling of the uterine collagen network.

## 2. Methods

### 2.1 Experimental inflation setup and microCT imaging

#### Tissue preparation

All animal experiments were approved by the Institutional Animal Care and Use Committee (IACUC) at the University of Illinois Urbana-Champaign and were performed in accordance with institutional guidelines and the Guide for the Care and Use of Laboratory Animals. Uteri were harvested from six, 6-month-old female CD-1 mice euthanized during diestrus. Both uterine horns were excised, and a 15-mm-long segment was isolated from each horn for inflation testing after removal of surrounding connective and adipose tissue. The untreated control and GA-treated groups each consisted of n = 6 uterine horn segments. Throughout specimen preparation, tissues were maintained submerged in Krebs buffer for hydration. Uterine horns assigned to the treatment group were immersed in 0.04% GA solution for 4 h at 4 °C. Following treatment, tissues were rinsed thoroughly in phosphate-buffered saline (PBS) to remove residual fixative. The GA concentration and exposure time were selected based on preliminary dose–response experiments. Higher concentrations produced excessive stiffening and altered the characteristic nonlinear mechanical response of the uterine wall.

#### Balloon catheter–based inflation system

To apply controlled, physiologically relevant internal loading to the murine uterus, a balloon catheter–based inflation system was employed using a 1.0 mm Sapphire II Pro semi-compliant balloon catheter (OrbusNeich, Hong Kong). This catheter was selected for its small diameter, flexibility, and controlled radial expansion characteristics, making it suitable for inflation of a soft hollow organ such as the mouse uterus.

The balloon conformed to the inner geometry of the uterus during inflation, providing an internal seal and approximately symmetric radial transmission of luminal pressure. This produced predominantly radial–circumferential deformation of the uterine wall and provided well-defined loading and boundary conditions for the finite element model.

#### Inflation protocol and pressure monitoring

Volume-controlled inflation experiments were performed using a dual-syringe pump (Fusion 200 dual-syringe pump, Chemyx Inc., Stafford, TX, USA), with one syringe connected to a balloon catheter mounted in the tissue and a second syringe connected to an identical catheter open to air as a reference. The two lines were coupled to a differential pressure sensor (Honeywell ABPMRRV015PD2A3; ±15 psi full-scale range, 0.25% full-scale accuracy), and pressure signals were acquired at 1 Hz using an Arduino microcontroller to measure intraluminal pressure relative to atmospheric pressure (Fig. 1A). This differential setup compensated for the pressure required to inflate the balloon itself, such that the recorded pressure primarily reflected tissue resistance. Fluid was infused at a constant rate of 0.02 mL/min using a 10 mL syringe to provide slow and controlled inflation. As the uterus could not remain submerged within the microCT system, Krebs buffer was applied to the tissue surface at approximately one-minute intervals to maintain hydration.

**Fig. 1.**
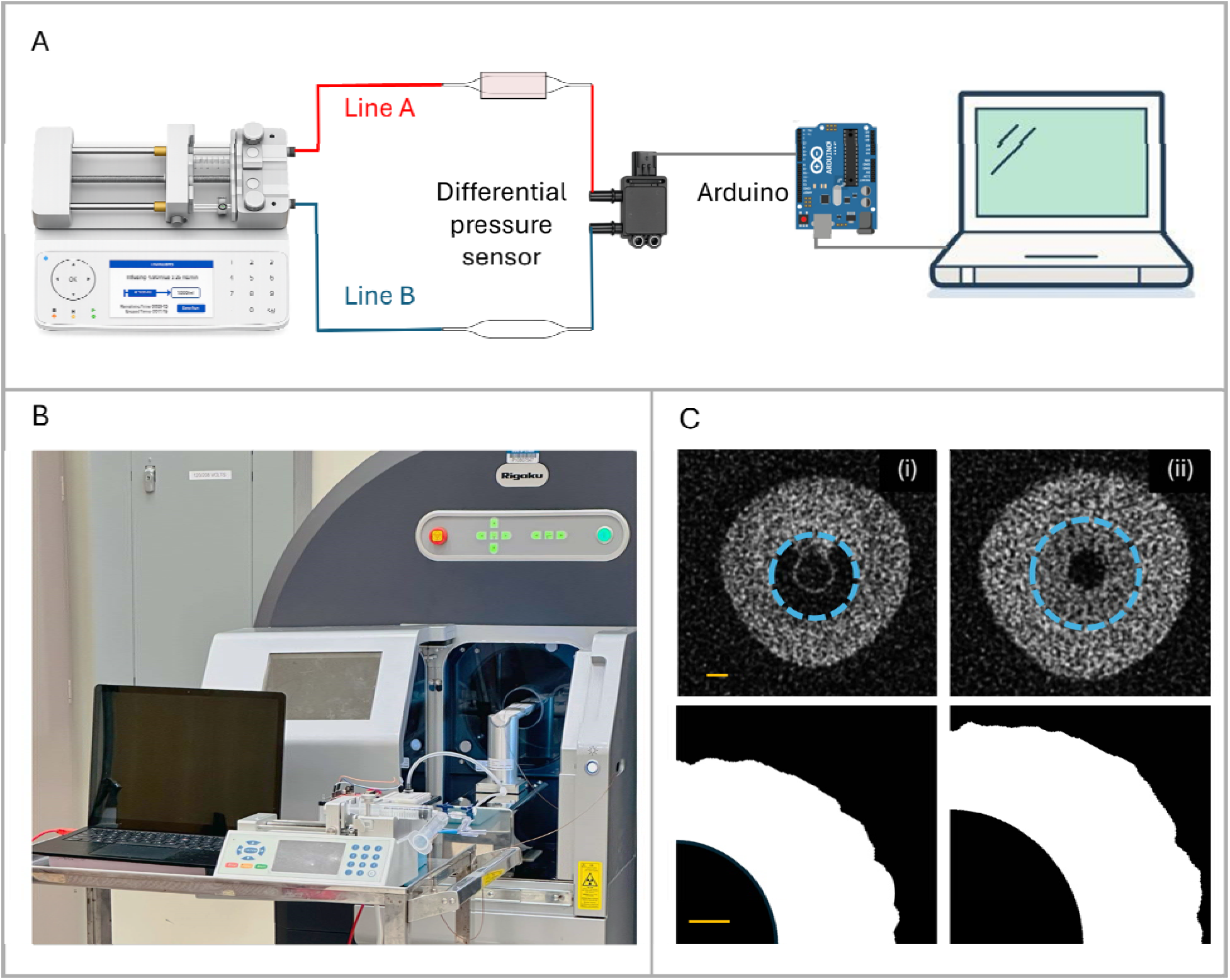
Experimental workflow for volume-controlled uterine inflation and image acquisition. (A) A schematic of the custom inflation system shows the syringe pump, fluid lines, differential pressure sensor, Arduino-based pressure acquisition, and computer interface. (B) This experimental setup was positioned in the microCT system to image the uterus during controlled inflation. (C) Representative microCT cross-sections of the murine uterus show images taken at two inflation states. Top panels show raw reconstructed slices; the cyan dashed line outlines the fluid-filled balloon lumen, which appears gray due to the presence of fluid, and bottom panels show the segmented representative uterine wall quadrant used for iFEA. Scale bars denote 250 μm

Prior to data collection, each specimen underwent three inflation–deflation preconditioning cycles to a maximum balloon volume of 0.20 mL to establish a repeatable mechanical response. Following preconditioning, the uterus was inflated in a stepwise manner by applying equal volumetric increments of 0.025 mL sequentially until a maximum balloon volume of 0.20 mL was reached. After each volume increment, fluid delivery was paused for approximately one minute to allow pressure stabilization prior to image acquisition. Following each inflation step, the measured tissue pressure reached a stable plateau within approximately 20 s (Supplementary Fig. S1E). Pressure–volume responses were subsequently quantified using inflation work (calculated as the area under the pressure–volume curve), linear stiffness between 0.10 and 0.20 mL inflation volume, and maximum inflation resistance at 0.20 mL.

#### Image acquisition

High-resolution microCT imaging was used to capture the three-dimensional geometry of the uterus at the reference (pre-inflation) state and at each subsequent inflation step using the Rigaku CTLab GX130 (Fig. 1B).

Scans were acquired with an isotropic voxel size of 10 µm, providing sufficient spatial resolution to resolve both the inner and outer boundaries of the uterine wall across inflation states. The scan field-of-view was 5×5×5mm, which encompassed the uterine segment mounted on the balloon catheter at all inflation levels.

A microCT scan was acquired at the reference state and after each of the eight sequential inflation increments, resulting in nine imaging states per specimen. Each scan required approximately four minutes to complete. Imaging was performed after each inflation step and the subsequent pressure stabilization period, allowing the tissue to reach a quasi-equilibrium configuration prior to image acquisition. This stepwise imaging approach enabled precise temporal alignment between applied volume, internal pressure, and tissue deformation, thereby allowing displacement estimation across the inflation steps.

#### Boundary segmentation and contour extraction

MicroCT images were processed using custom Python scripts to determine uterine wall boundaries at each inflation state. Representative raw microCT images and the corresponding segmented uterine wall quadrant are shown in Fig. 1C (top and bottom panels, respectively). Raw grayscale images were preprocessed using Gaussian blurring to reduce noise while preserving boundary edges. Intensity thresholding was then applied to isolate uterine tissue from the surrounding background and balloon, producing binary masks for contour extraction. The inner and outer uterine wall boundaries were subsequently identified from the binary masks using an active contour (snake) algorithm [47, 48], which generated smooth, closed representations of the tissue–balloon and external tissue interfaces. The resulting boundary coordinates were interpolated to obtain continuous contour representations at each inflation state. Deformation was then quantified by comparing corresponding boundary geometries between inflation states.

#### Displacement computation

For each inflation step, inner and outer boundary contours were expressed in a polar coordinate system centered at the lumen, and boundary locations were sampled at uniform angular increments of 0.5°. Corresponding boundary points between the reference and deformed configurations were identified based on their shared angular position. All coordinate measurements were converted from pixels to millimeters using the known isotropic voxel size of the scan.

### 2.2 Finite Element Model

#### Geometry reconstruction

Finite element geometries of the uterine wall were reconstructed directly from experimentally measured inner and outer boundary contours obtained from microCT. For each sample, the extracted two-dimensional boundary point clouds were imported into FreeCAD, where they were converted into closed splines and used to generate a solid representation of the uterine cross-section. The resulting geometry was then exported as a STEP file and imported into Abaqus for meshing and finite element analysis. This workflow enabled the construction of sample-specific geometries that closely reflected the experimentally observed anatomy.

The uterine wall was represented as a thick-walled deformable pressure vessel modeled using experimentally measured inner and outer boundaries. Although the uterus comprises multiple layers (endometrium, circumferential myometrium, and longitudinal myometrium), these layers were not modeled separately; instead, the uterus was modeled as a homogeneous continuum with two fiber families: circumferential and longitudinal. This simplified representation provides a balance between physiological relevance and parameter identifiability, enabling robust estimation of constitutive parameters from experimentally measured boundary displacements.

#### Constitutive model

The mechanical behavior of the uterine wall was modeled using the GOH hyperelastic constitutive formulation, which is well-suited for fiber-reinforced soft tissues exhibiting anisotropic and nonlinear mechanical behavior. The total strain energy density function can be expressed as:

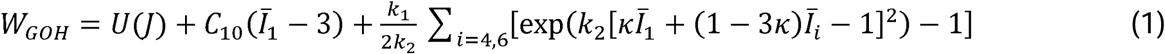

where the volumetric contribution is given by

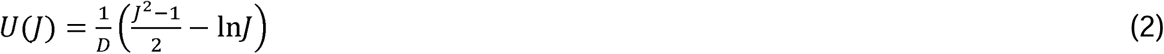

*J*=det(**F**) is the volumetric Jacobian, where **F** is the deformation gradient, and **D** governs material compressibility. As *D*→0, the material approaches incompressible behavior.*C*_10_ represents the stiffness of the non-collagenous ground matrix and primarily governs the tissue response in shear and at low strains. The parameter *k*_1_ controls the stiffness of the collagen fiber phase and dictates the magnitude or density of fiber reinforcement. The parameter *k*_2_ governs the nonlinearity of the fiber response and is associated with gradual fiber recruitment during stretching. Fiber dispersion is characterized by *k*, which quantifies the spread of fiber orientations about a preferred direction, with *k* = 0 corresponding to perfectly aligned fibers and *k* = 1/3 representing an isotropic distribution. *Ī*_1_ is the first isochoric invariant, while *Ī*_4_ and *Ī*_6_ represent the squared stretches along the two fiber families. The formulation assumes two symmetrically arranged collagen fiber families.

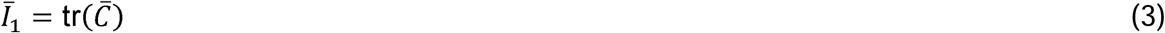

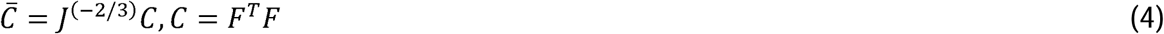

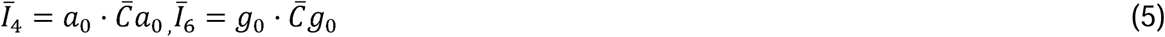

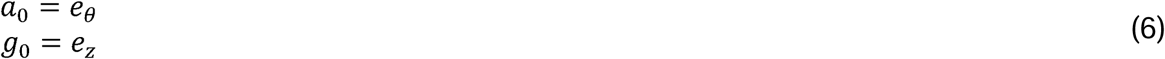

where *e_θ_* and *e_z_* denote the circumferential and longitudinal unit vectors.

In the present study, the uterine tissue was assumed to be incompressible (*J* ≈ 1). Accordingly, hybrid finite elements available in Abaqus were employed, in which hydrostatic pressure is introduced as an independent field variable and incompressibility is enforced through a mixed displacement-pressure formulation. Under this formulation, Eq. (1) can be equivalently written as:

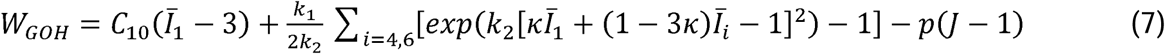

Where *p* is the hydrostatic pressure acting as a Lagrange multiplier that enforces the incompressibility constraint.

### Balloon material model

The mechanical behavior of the balloon catheter was explicitly incorporated into the finite element model to accurately represent tissue–balloon interaction during inflation. The balloon was modeled using a 4-term isotropic Ogden hyperelastic formulation [30, 49, 50],

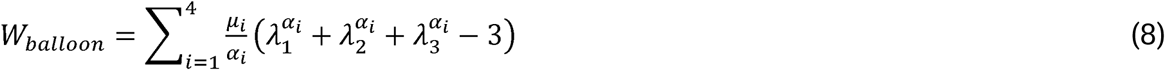

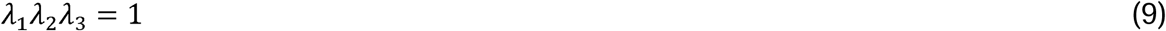

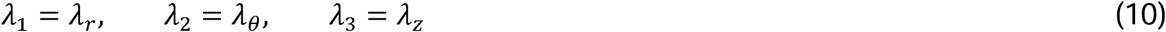

To characterize the balloon material independently of the uterine tissue, balloon-only inflation experiments were performed and imaged using Zeiss Xradia Versa 360 nanoscale-resolution microCT scanner. Pressure–diameter measurements extracted from these scans were used to calibrate the Ogden parameters using the same inverse finite element framework employed for uterine parameter estimation. Material parameters were identified by minimizing the objective function *Φ_balloon_*

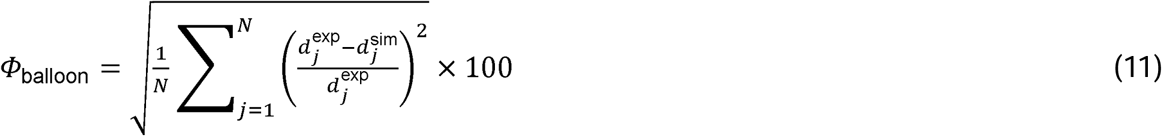

where 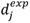 and 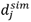 represent the experimental and simulated balloon diameters at the *j^th^* loading increment and *N* is the total number of measurement points.

Representative optimization histories are shown in Supplementary Fig. S2, and the optimized Ogden parameters are summarized in Table 1.

**Table 1.** Optimized four-term Ogden material parameters obtained from inverse finite element calibration of the balloon using balloon-only inflation experiments.

| Parameter | Value |
| --- | --- |
| $\mu_1$ | 0.035 |
| $\alpha_1$ | 0.591 |
| $\mu_2$ | 0.11 |
| $\alpha_2$ | 2.705 |
| $\mu_3$ | 4.267 |
| $\alpha_3$ | 8.861 |
| $\mu_4$ | 0.23 |
| $\alpha_4$ | -2.738 |

The balloon exhibited repeatable inflation behavior with minimal dependence on inflation rate, and the calibrated model accurately reproduced the experimental balloon response (Supplementary Fig. S1B-D). These results indicate that the balloon provided a stable and reproducible loading interface throughout the inflation experiments.

Contact between the balloon and uterine wall was modeled using a low Coulomb friction coefficient (*μ_f_* = 0.03), allowing relative sliding with minimal resistance and approximating the glycerol-lubricated experimental environment [51, 52]. A friction sensitivity analysis confirmed that the results were insensitive to the assumed friction coefficient (Supplementary Fig. S3).

#### Representative quadrant selection

The uterine cross-section has non-uniform wall thickness around the circumference. As the iFEA was performed on a representative quarter of the uterine cross-section, using a randomly selected region of the uterus may bias parameter estimation toward locally thick or thin regions that are not representative of the overall tissue structure. Although modeling the full 360° uterine cross-section would capture circumferential heterogeneity, such simulations substantially increase the computational cost of iFEA. Therefore, a representative 90° sector was selected to reduce computational expense while preserving wall-thickness characteristics representative of the overall cross-section.

For each sector, local wall thickness was computed at uniformly spaced angular positions as the radial distance between the inner and outer boundary contours. Specifically, at angular position *θ_i_*,

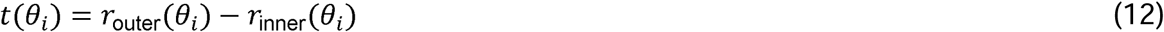

Where, *r_outer_* and *r_inner_* denote the radial distances from the lumen center to the inner and outer boundaries, respectively.

Wall thickness was evaluated at 1° increments, yielding 360 circumferential thickness measurements. For each candidate 90° sector beginning at angular index *k*, the mean sector thickness was calculated as

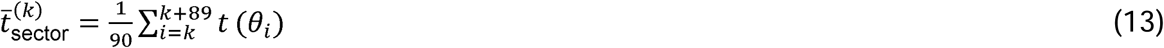

The global mean wall thickness across the entire cross-section was similarly computed as

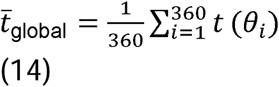

The representative quadrant was then identified as the 90° sector whose mean wall thickness was closest to the global mean thickness according to

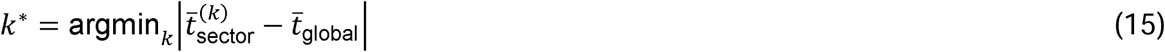

The representative quadrant was subsequently defined as the sector beginning at index *k*\* This procedure selected the 90° sector whose mean wall thickness best represented that of the complete uterine cross-section while substantially reducing the computational cost of iFEA (Supplementary Fig. S4).

### Boundary conditions and loading

The selected representative uterine wall quadrant was modeled using a quarter-symmetry finite element representation (Fig. 2A). Symmetry boundary conditions were imposed on two orthogonal planar faces of the model. On one symmetry plane, displacement in the global X-direction was constrained (*u_x_*=0), while on the second orthogonal plane, orthogonal plane, displacement in the global Y-direction was constrained (*u_y_*=0). These constraints correspond to symmetry about the YZ and XZ planes, respectively. Displacements tangent to each symmetry plane were left unconstrained, allowing in-plane deformation during inflation. In addition, displacement normal to the longitudinal end faces was constrained (*u_z_*=0), approximating plane-strain condition over the imaged segment.

**Fig. 2.**
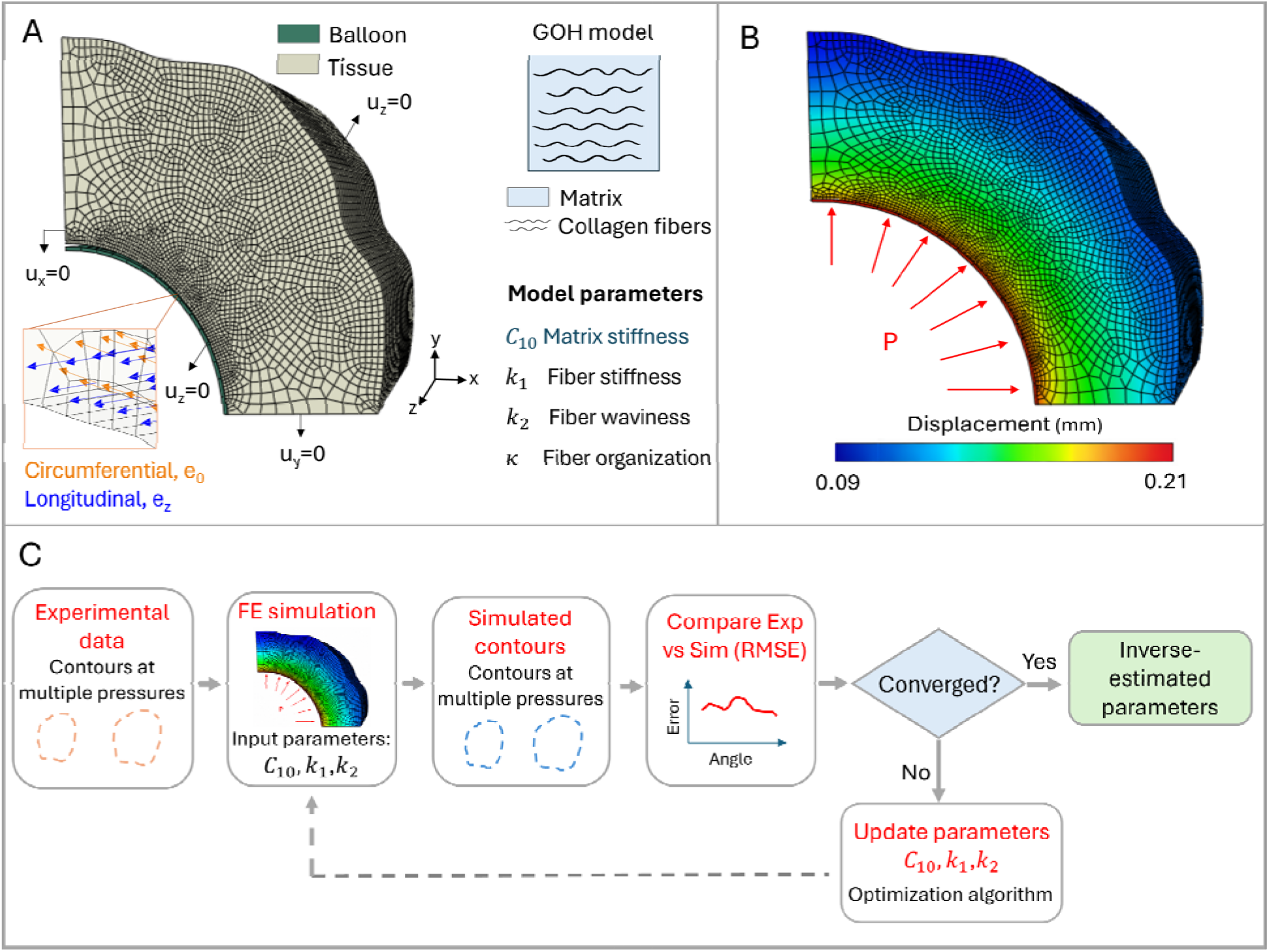
Image-informed inverse finite element analysis (iFEA) framework for constitutive parameter estimation from uterine inflation experiments. (A) A representative finite element mesh of the uterine quadrant model shown with applied displacement constraints and internal pressure loading. The constitutive behavior of the tissue was modeled using the GOH formulation, which accounts for the isotropic matrix and embedded collagen fiber families. Circumferential and longitudinal fiber orientations were incorporated to represent uterine collagen organization. (B) An example finite element simulation is shown with the distribution in displacement magnitude during loading. (C) Schematic illustrating the iFEA workflow. Experimental contours extracted from microCT images were compared with simulated contours generated from the Finite Element (FE) model. The optimization algorithm iteratively updated the GOH parameters (*C*_10_, *k*_1_ and *k*_2_) until convergence

The pressure prescribed to the balloon in the model was calculated as the sum of the experimentally measured tissue pressure and the balloon pressure associated with the same inflation volume, as determined from the balloon-only pressure–volume curve. This total pressure was transmitted to the uterine wall through explicit balloon–tissue contact, consistent with previous computational models of balloon expansion in soft tissues [30]. A representative displacement field obtained from the finite element model is shown in Fig. 2B.

### Mesh convergence analysis

Mesh convergence was evaluated by comparing contour predictions obtained using coarse, medium, and fine meshes. Root mean square error (RMSE) relative to the fine-mesh solution was computed separately for the inner and outer boundaries, and computational time was recorded for each mesh density. Based on the balance between accuracy and computational cost, the medium mesh was selected for all subsequent simulations (Supplementary Fig. S5).

### 2.3 Inverse Finite Element Analysis (iFEA)

#### Objective function

iFEA was performed by minimizing the mismatch between experimentally measured and computationally predicted uterine boundary geometries (Fig. 2C). The objective function, *Φ*, was formulated using radial profiles extracted from the inner and outer uterine boundaries, enabling direct comparison between simulated and experimentally measured boundary geometries at each inflation state. The objective function was defined as:

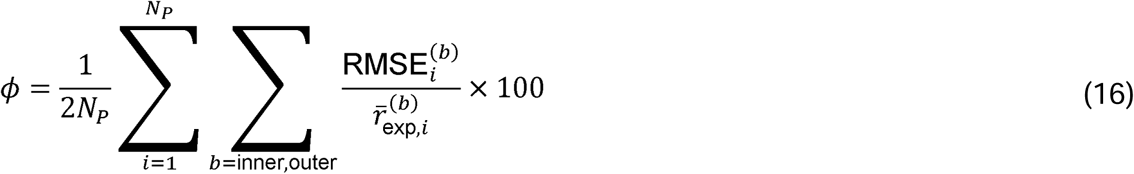

where the boundary error for each inflation state was quantified using the RMSE,

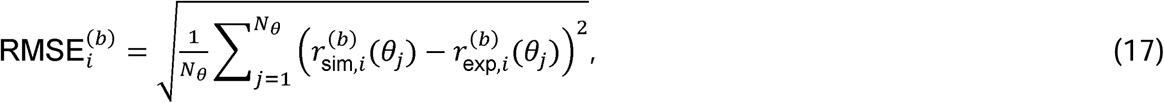

and

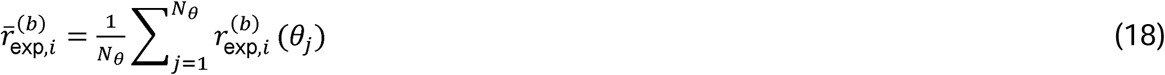

Here, 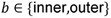 denotes the boundary, *i* = 1,2…….., *N_p_* indexes the inflation states, *j* = 1,2…..,*N_θ_* indexes the angular sampling locations, and 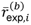 is the mean experimental boundary radius at inflation state i

The objective function was evaluated using four discrete inflation states for each specimen. These states were selected from the experimentally measured pressure– volume curves (Fig. 3A) to capture key regions of the inflation response while limiting computational cost. For each specimen, the selected states included a low-pressure point within the initial toe region, two pressure points surrounding the inflection region, and a higher-pressure point corresponding to a later stage of inflation. The optimization points used for contour matching are highlighted in Fig. 3A. Because untreated and GA-treated tissues exhibited different pressure–volume responses, different inflation states were selected for each group to represent comparable stages of inflation.

**Fig. 3.**
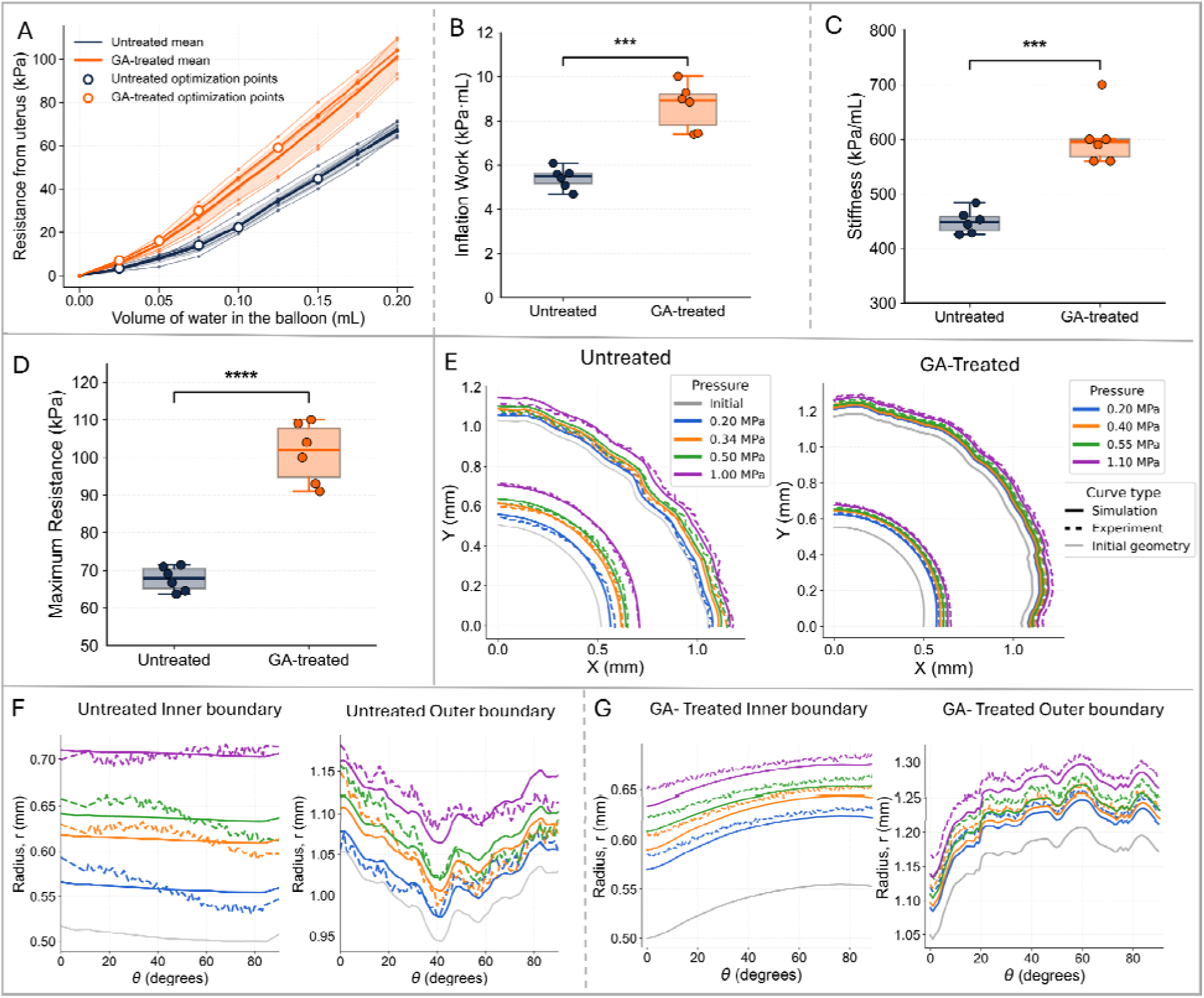
Experimental inflation behavior and iFEA model validation for untreated and GA-treated uterine tissues. (A) Pressure–volume responses during balloon inflation differed for untreated and GA-treated samples. Thin lines represent individual samples, and thick lines represent group means. Open circles indicate the inflation states selected for contour matching and iFEA. GA-treated tissues exhibited higher resistance to inflation compared to untreated tissues across the loading range. (B–D) Quantitative metrics summarizing inflation behavior, including inflation work (B), linear stiffness (C), and maximum resistance (D) (***p < 0.001, ****p < 0.0001). (E) Representative contour comparisons between experimental and simulated deformations for untreated and GA-treated tissues at multiple total applied pressure levels. The pressure values correspond to the total pressure applied in the finite element model, including both balloon and tissue contributions. Solid lines indicate simulated contours, dashed lines indicate experimental contours, and gray lines represent the initial geometry. (F–G) Radius– angle ( – ) profiles of the inner and outer boundaries for representative untreated and GA-treated samples, respectively, demonstrate good agreement between experimental and simulated contours across all pressure levels for both groups. Line styles are identical to those shown in (E)

**Fig. 4.**
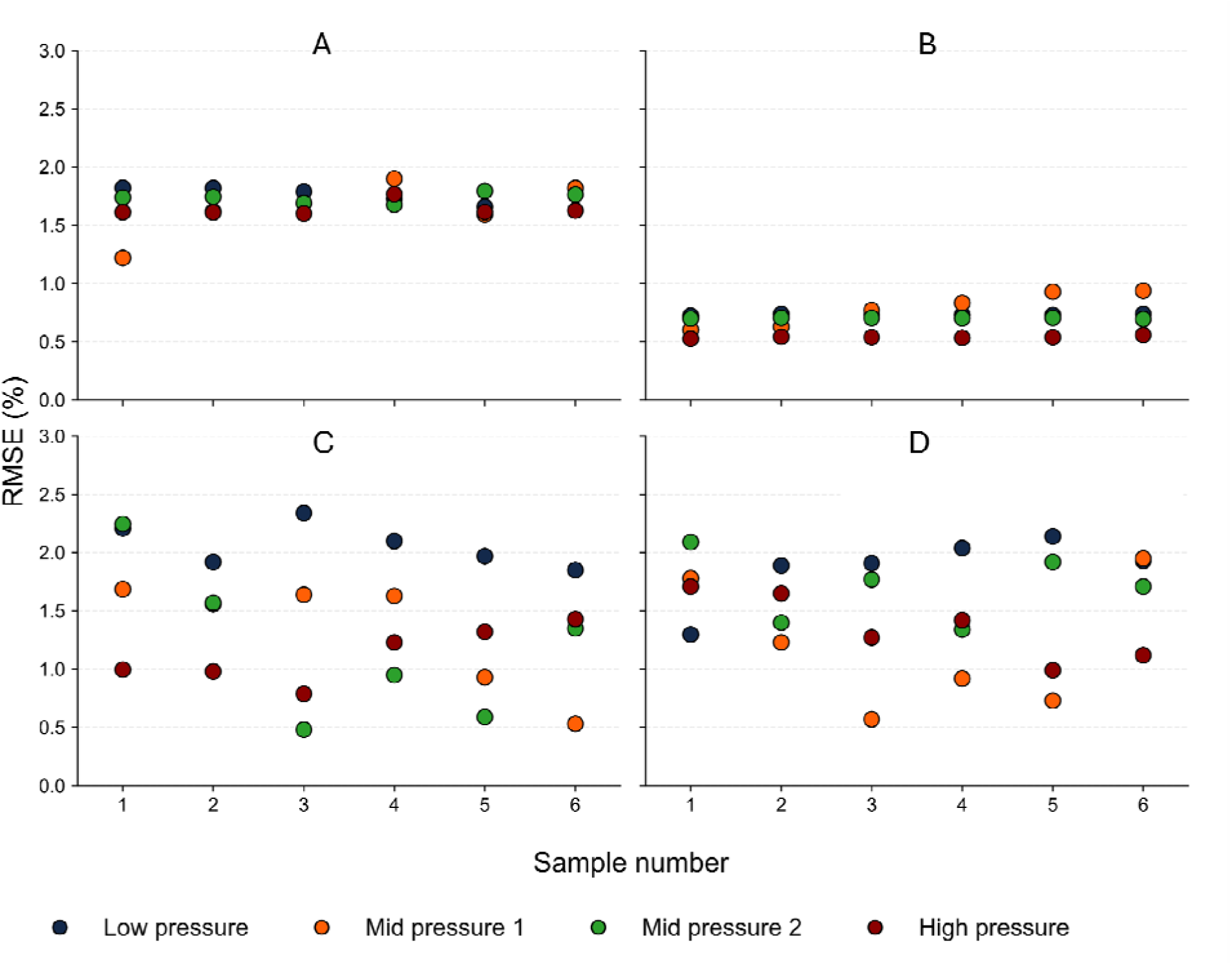
Root mean square error (RMSE) between experimentally measured and simulated boundaries for untreated and GA-treated uterine tissues. Panels show the RMSE for the (A) untreated inner boundary, (B) untreated outer boundary, (C) GA-treated inner boundary, and (D) GA-treated outer boundary. Colors indicate the four inflation states used during inverse optimization (low pressure, mid pressure 1, mid pressure 2, and high pressure). RMSE values remained below 3% for all samples and inflation states

#### Optimization strategy

Constitutive parameters *C*_10_, *k*_1_ and *k*_2_ were estimated using iFEA. The fiber dispersion parameter *k* was fixed at 0.1 throughout the optimization and was not treated as an optimization variable. The selected value was derived from previously reported image-based spherical variance measurements and weighted according to the relative collagen contributions of the uterine layers to obtain a representative tissue-level dispersion parameter [9]. Fixing *k* reduced parameter coupling during optimization and improved the identifiability of the remaining constitutive parameters.

Parameters were estimated using a two-stage optimization strategy consisting of a global optimization stage followed by a local optimization stage. A differential evolution (DE) algorithm was initially employed to explore the parameter space and identify a suitable starting region for subsequent optimization [53]. Sample-specific parameter estimation was then performed using the Nelder–Mead simplex algorithm [54]. For each candidate parameter set, a forward finite element simulation was performed, and the predicted boundary radial profiles were compared with the experimentally measured profiles using the objective function defined in Eq. (16). Optimization was terminated when the relative change in the objective function was less than 1% over five consecutive iterations. A representative optimization history is shown in Supplementary Fig. S6. To limit computational expense, a maximum of 50 Nelder–Mead iterations was imposed.

#### Reconstruction of constitutive response

To facilitate interpretation of the estimated constitutive parameters, forward finite element simulations were performed using the optimized material parameters (Fig. 5C). Circumferential stress and logarithmic strain were extracted from representative elements within the uterine wall over the inflation loading range. The resulting stress-strain responses were used to compare nonlinear mechanical behavior between untreated and GA-treated tissues and to identify differences in the toe and strain-stiffening regions.

**Fig. 5.**
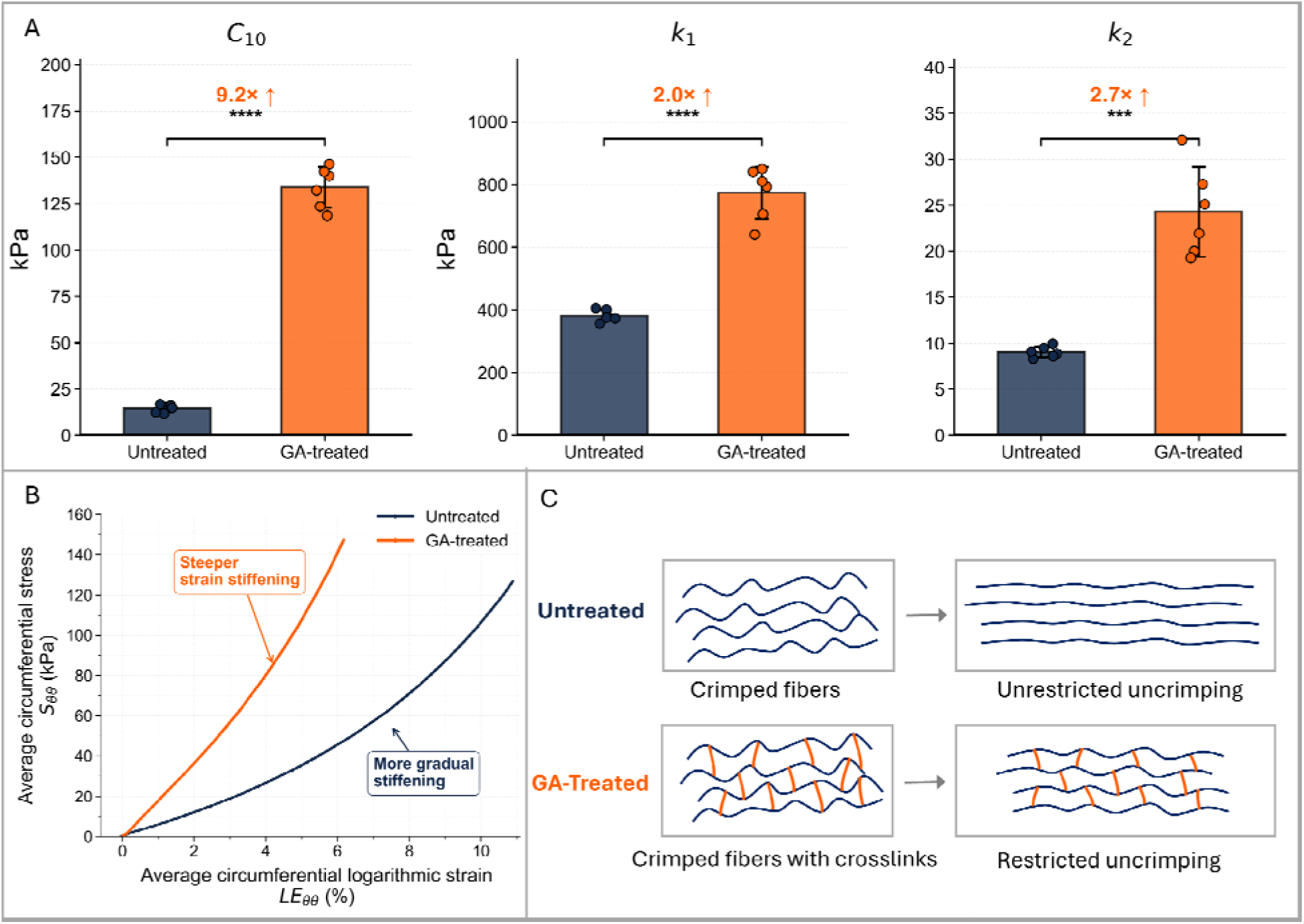
GA treatment altered the estimated constitutive parameters and the resulting mechanical response of the uterine wall. (A) GA treatment significantly increased the GOH model parameters and Bars represent mean ± standard deviation, points represent individual samples, and fold changes relative to untreated tissues are indicated above each comparison. Statistical significance was determined using Welch’s t-test ( , ). (B) Reconstructed circumferential stress–strain response generated using the mean constitutive parameter estimates illustrates the altered mechanical behavior following GA treatment. GA-treated tissues exhibited an earlier onset of strain stiffening and a steeper increase in stiffness with deformation, whereas untreated tissues displayed a broader toe region and a more gradual stiffening response. (C) Conceptual schematic illustrating the proposed effect of GA crosslinking on collagen recruitment. In untreated tissue, collagen fibers undergo progressive uncrimping during loading, whereas GA-induced crosslinks are proposed to restrict fiber uncrimping, leading to earlier mechanical engagement during deformation

### 2.4 Sensitivity and robustness analysis

To assess the robustness of the optimized constitutive parameters and evaluate the sensitivity of model predictions to parameter variations, sensitivity analysis was performed following iFEA optimization (Fig. 6). Because constitutive parameter estimation may yield multiple parameter combinations that produce similarly accurate model predictions, evaluating parameter sensitivity and identifiability is important for interpreting optimized material properties [55–60]. This analysis was designed to determine whether small perturbations in material parameters lead to disproportionate changes in model response and to characterize the local structure of the error landscape around the optimized solution.

**Fig. 6.**
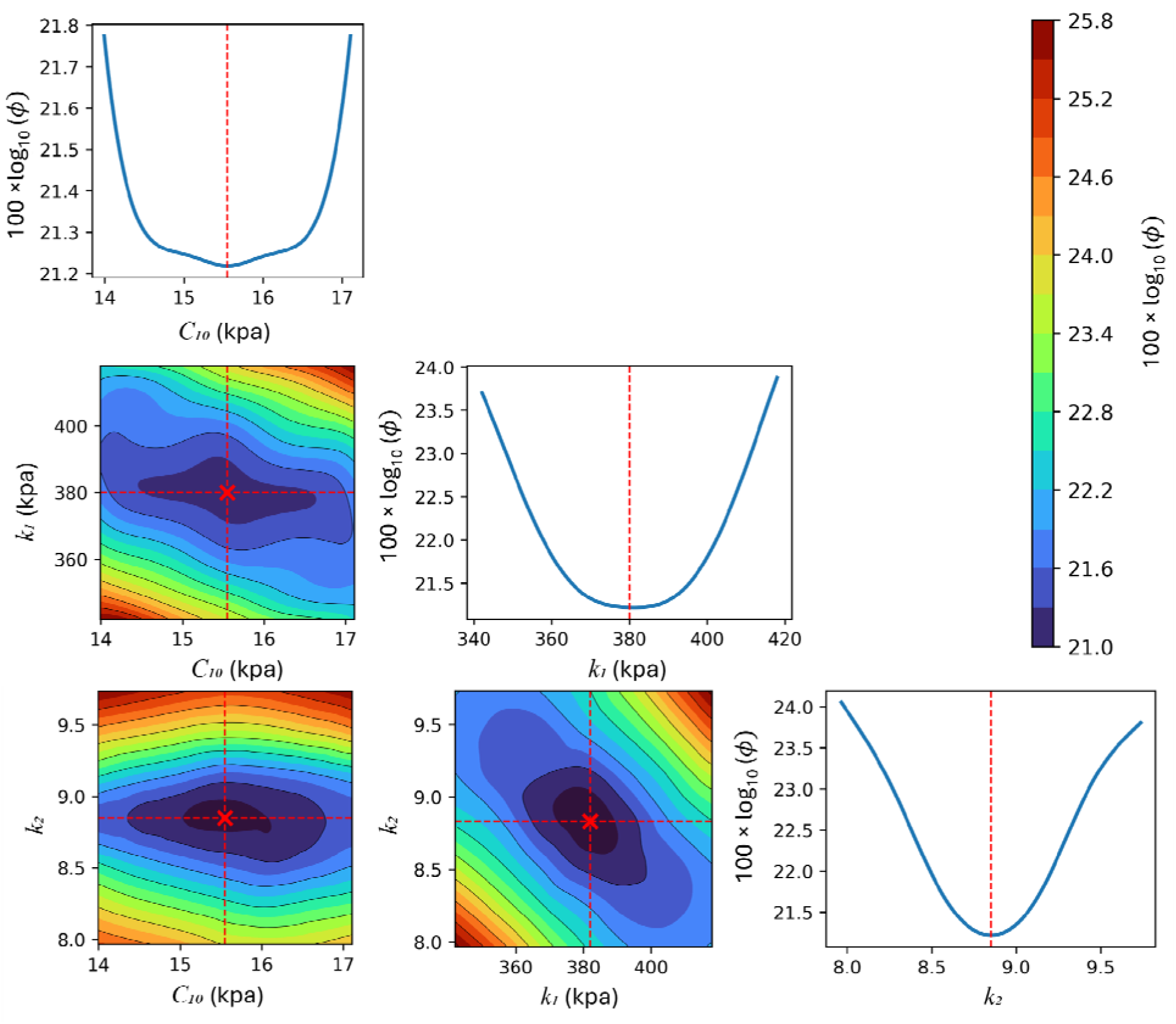
One- and two-parameter objective function plots demonstrate a unique minimum. Diagonal panels show one-parameter sensitivity plots of the objective function in terms of individual material parameters C_10_, k_1_, and k_2_, with all other parameters held fixed at their optimized values. Off-diagonal panels show two-parameter objective function surfaces for selected parameter pairs (C₁₀–k₁, C₁₀–k₂, and k₁–k₂). Red markers indicate the location of the minimum error within the explored parameter space. The color bar indicates the magnitude of log_10_( )

Grid-based sensitivity analyses were performed around the best-fit solution. The analysis was performed for a representative untreated sample using its optimized parameter set. One-parameter sensitivity analyses were conducted by perturbing each material parameter individually while holding all remaining parameters fixed at their optimized values, allowing evaluation of the local error response along each parameter dimension. Two-parameter sweeps were conducted by varying selected parameter pairs around optimized values, using a 7×7 grid spanning a ±10% range for each parameter. For each parameter combination, forward finite element simulations were performed, and the resulting boundary radial profiles were compared with experimental measurements by evaluating the same objective function employed during optimization.

To further quantify parameter identifiability and characterize the local geometry of the objective-function landscape, curvature-based metrics were computed from the two-parameter sensitivity maps. For each parameter pair, the objective function was evaluated on a regular grid surrounding the optimized solution, and a local Hessian matrix was estimated using finite-difference approximations. Eigenvalues of the Hessian were used to quantify local curvature near the optimized solution. The minimum eigenvalue (λ_min_) was used as a measure of the weakest constrained direction in parameter space, with larger values indicating stronger local identifiability. Basin anisotropy was quantified using the Hessian condition number, defined as the ratio of the largest to smallest eigenvalue magnitudes. Lower condition numbers indicate a more isotropic objective basin, whereas larger values indicate an elongated valley.

### 2.5 Statistical analysis

Estimated constitutive parameters obtained from iFEA were compared between the untreated and GA-treated groups. Data are reported as mean ± standard deviation. Statistical differences between groups were evaluated using Welch’s two-sample t-test to account for potential differences in variance. Statistical significance was defined as p < 0.05.

## Results

### 3.1 Inflation response of untreated and GA-treated tissues

Both untreated and GA-treated tissues exhibited nonlinear pressure-volume behavior characterized by progressive stiffening with increasing inflation volume (Fig. 3A). Across the entire loading range, GA-treated tissues consistently exhibited greater resistance to inflation than untreated tissues, indicating an overall increase in tissue stiffness following treatment. Although some inter-sample variability was present within each group, clear separation between the two groups was maintained at all inflation volumes.

To quantify these differences, inflation work, linear stiffness, and maximum resistance were computed from the pressure–volume responses (Fig. 3B–D). GA treatment significantly increased inflation work, linear stiffness, and maximum resistance. Together, these results demonstrate that GA treatment significantly increased the mechanical resistance of the uterine wall during inflation.

### 3.2 Finite element reproduction of experimental deformation

Representative comparisons between experimentally measured and simulated contour deformations are shown in Fig. 3E–G for untreated and GA-treated tissues. Across all pressure levels, the finite element model reproduced both inner and outer boundary deformations, demonstrating close agreement between simulations and experimental observations.

Contour overlays revealed that the model successfully captured both the overall circumferential expansion of the uterus and local variations associated with nonuniform wall geometry. Similar agreement was observed for untreated and GA-treated specimens, indicating that the framework remained valid despite the changes in tissue stiffness induced by crosslinking. Radius–angle profiles further demonstrated close correspondence between experimental and simulated contours across all inflation states (Fig. 3F-G).

To evaluate model performance across all samples, errors were quantified using the RMSE between experimental and simulated boundaries (Fig. 4). Untreated tissues exhibited consistently low RMSE values with relatively little variation across samples and pressure levels. GA-treated tissues showed a broader distribution of RMSE values, particularly for the inner boundary, although errors remained low overall. Across both groups, RMSE values remained below 3%, confirming that the inverse framework accurately captures deformation behavior over a wide range of mechanical responses. Complete contour comparisons for all specimens are provided in Supplementary Fig. S7.

### 3.3 Effect of GA treatment on constitutive parameters

Optimized GOH material parameters for untreated and GA-treated tissues are summarized in Fig. 5A. Across untreated tissues, the optimized parameter values were highly consistent between samples. The matrix stiffness parameter *C*_10_ averaged 14.6 ± 2.1 kPa (CV = 14.5%), while the fiber stiffness parameter *k*_1_ averaged 381.9 ± 18.3 kPa (CV = 4.8%). The fiber recruitment parameter *k*_2_ averaged 9.1 ± 0.6 (CV = 6.7%). All parameters varied by less than 15% across untreated samples, indicating that the inverse framework produced reproducible constitutive parameter estimates.

The same inverse analysis was performed on GA-treated samples. GA treatment produced significant increases in all three constitutive parameters (Fig. 5A). The matrix stiffness parameter *C*_10_ increased from 14.6 kPa to 133.8 kPa (9.2-fold increase; CV = 8.3%). Similarly, *k*_1_ increased from 381.9 kPa to 773.2 kPa (2.0-fold increase; CV = 10.8%), while *k*_2_ increased from 9.1 to 24.3 (2.7-fold increase; CV = 20.1%). These parameter changes indicate a significantly stiffer tissue response following treatment and are consistent with the enhanced strain-stiffening behavior observed in the simulated stress–strain response (Fig. 5B).

To illustrate the mechanical implications of these parameter changes, representative stress–strain curves were generated using the mean parameter sets for untreated and GA-treated tissues (Fig. 5B). Untreated tissues exhibited a broad toe region followed by a gradual increase in stiffness with strain. In contrast, GA-treated tissues showed a markedly steeper response and exhibited strain stiffening at lower strains. The steeper response observed in GA-treated tissues is consistent with earlier mechanical engagement of collagen fibers, resulting in a more rapid increase in tissue stiffness during inflation. Figure 5C provides a conceptual schematic of the proposed collagen recruitment behavior associated with these parameter changes.

### 3.4 Sensitivity and robustness of inverse solutions

One-parameter perturbation analyses (Fig. 6, diagonal panels) revealed unimodal and symmetric error profiles for and . In all cases, the objective function exhibited a single minimum at the optimized parameter value and increased smoothly as parameters were perturbed away from the optimized solution. No secondary minima was observed within the explored parameter ranges, supporting the existence of a unique local minimum.

To quantitatively characterize parameter identifiability, curvature-based metrics were computed from the local Hessian of the objective function. The *k*_1_-*k*_2_ parameter pair exhibited the strongest local parameter constraint, with the largest minimum eigenvalue (*λ_min_*=25.8) and the lowest condition number (2.02), indicating a well-defined and relatively isotropic objective basin. In contrast, the *C*_10_-*k*_1_ pair exhibited the smallest minimum eigenvalue (*λ_min_*=2.69), indicating weaker local parameter constraint, while the *C*_10_-*k*_2_ pair showed the highest condition number (6.27), reflecting the greatest degree of parameter coupling.

Consistent with these quantitative metrics, the two-parameter sensitivity maps (Fig. 6) exhibited a single basin of minimum error for all parameter combinations. Although the degree of curvature and anisotropy varied among parameter pairs, all surfaces converged to a unique minimum within the explored range, supporting the practical identifiability of the iFEA solution.

## 4. Discussion

### 4.1 Inflation behavior and constitutive framework

This study presents a combined balloon catheter–based inflation experiment and iFEA framework for estimating constitutive parameters of the murine uterine wall from microCT-derived deformation data. By integrating experimentally measured displacement contours with image-informed finite element modeling, the proposed framework enables quantitative characterization of uterine mechanical behavior and provides mechanistic insight into how tissue crosslinking alters the constitutive response.

Across all specimens, inflation experiments showed a nonlinear pressure–volume response characterized by an initially compliant region followed by progressive stiffening at higher inflation volumes (Fig. 3A). This nonlinear behavior is characteristic of many collagen-rich soft tissues and reflects the increasing contribution of collagen fibers to load bearing with increasing strain. At lower strains, the isotropic contribution dominates the mechanical response, whereas the fiber contribution becomes progressively more important with increasing deformation. The repeatability of the pressure–volume response demonstrates that the experimental setup provides consistent loading conditions and forms a reliable basis for inverse parameter estimation.

The observed inflation behavior aligns well with the formulation of the GOH constitutive model, which explicitly separates isotropic matrix contributions from anisotropic fiber recruitment. In this formulation, early inflation is dominated by the isotropic contribution represented by *C*_10,_ whereas increasing inflation progressively engages the fiber families governed by *k*_1_ and *k*_2_. The ability of the finite element model to reproduce experimentally measured deformations with low contour-matching error across all inflation states supports that this constitutive framework is reasonable for describing the passive mechanical behavior of the uterine wall under inflation (Fig. 3E–G).

### 4.2 Constitutive effects of collagen crosslinking

A major finding of this study was the substantial shift in constitutive behavior following GA treatment. Compared with untreated tissues, GA-treated samples exhibited significantly higher resistance to inflation (Fig. 3A–D) and elevated values of and (Fig. 5A). Within the GOH framework, scales the magnitude of the fiber stress contribution and therefore reflects the overall load-bearing capacity of the collagen network, whereas *k_2_* governs the rate at which the fiber contribution increases with strain. The observed increases in both parameters indicate that GA treatment altered both the magnitude of the collagen-driven mechanical response and the manner in which collagen fibers became mechanically engaged during deformation. Consistent with this interpretation, reconstructed stress–strain responses exhibited increased stiffness and more rapid strain stiffening following treatment (Fig. 5B).

The proposed mechanistic interpretation of these changes is illustrated in Fig. 5C. GA forms covalent intermolecular crosslinks within collagen-rich tissues, increasing resistance to molecular sliding and restricting fibrillar mobility. In untreated tissue, collagen fibers progressively become load-bearing through mechanisms such as uncrimping, reorientation, and alignment with the loading direction. Following GA treatment, crosslinks may restrict these deformation mechanisms and promote load transfer to the collagen network at lower strains. Consequently, the increases in *k*_1_ and *k*_2_ are consistent with a collagen network that is both stiffer and more rapidly recruited during deformation. This pattern of GA-induced stiffening at the whole-tissue level is consistent with reports in other collagenous tissues [61–63]

Whereas the increases in *k*_1_ and *k*_2_ are readily interpreted within the classical GOH framework, the substantially larger increase observed in *C*_10_ requires additional consideration. In the classical GOH formulation, *C*_10_ represents the isotropic matrix contribution and therefore would not be expected to increase solely because of collagen crosslinking. However, because pressure in this experiment is applied to the luminal surface, deformation initiates in the endometrium and propagates outward through the uterine wall (Fig. 2A). Although the endometrium contains collagen at a lower concentration and with less preferential organization than the myometrium [9], its collagen network likely contributes to the early radial–circumferential (*r-θ*) shear resistance. In a loosely organized network, shear is accommodated primarily through fibril sliding and reorientation, whereas GA crosslinking limits this sliding by tethering adjacent fibrils together and forcing the network to resist shear more directly [61]. Therefore, *C*_10_ in the present study should be interpreted more broadly as an effective parameter that captures the early inflation resistance of the uterine wall, including contributions from the crosslinked endometrium, rather than strictly as a measure of non-collagenous matrix stiffness.

### 4.3 Identifiability, Limitations, and Future Directions

Inverse analysis yielded consistent constitutive parameter estimates across untreated specimens. The relatively small variability observed in *k*_1_ and *k*_2_ indicates that the fiberrelated mechanical response of the uterine wall was well constrained by the inflation data. Under inflation, circumferential collagen fibers undergo substantial stretch and 25 progressively contribute to load bearing capacity, allowing both fiber stiffness (*k*_1_) and nonlinear recruitment (*k*_2_) to be robustly identified from the measured boundary kinematics despite the simplified homogeneous tissue representation. In contrast, *C*_10_ exhibited greater variability than the fiber-related parameters (Fig. 5A). This likely reflects a combination of biological heterogeneity and the greater sensitivity of *C*_10_ to internal deformation mechanisms that cannot be independently resolved from boundary deformation measurements, including radial–circumferential (*r*-*θ*) shear

The sensitivity analyses support the identifiability of the inverse solutions (Fig. 6). Although moderate coupling was observed between some parameters, all error landscapes exhibited a single well-defined minimum, indicating stable optimization results. Even for the least identifiable parameter pair, the objective landscape remained well behaved and exhibited a distinct minimum. While the Hessian analysis characterizes only the local curvature of the objective function, the initial differential evolution stage performed a global search of the parameter space before local refinement, reducing the likelihood of convergence to a suboptimal local minimum.

The inflation response was most sensitive to the fiber-related parameters, suggesting that collagen fiber mechanics play the dominant role in governing uterine inflation behavior. These findings support the interpretation of the treatment-induced changes in material parameters as meaningful mechanical differences between groups. Furthermore, since the balloon material model was independently calibrated and held fixed throughout all inverse analyses, these results provide confidence that the predicted deformation remained governed by uterine constitutive behavior rather than by the balloon mechanics.

GA treatment was used as a large, controlled perturbation of tissue mechanics to establish proof-of-concept sensitivity to the proposed inverse framework. The framework successfully detected the major constitutive changes induced by collagen crosslinking. However, the smallest mechanical changes that can be detected remain unknown. Future studies using varying levels of mechanical perturbations and environmentally induced tissue remodeling will help define the detection limits and resolution of the proposed approach.

Several simplifying assumptions were made to enable this inverse analysis. The uterine wall was modeled as a homogeneous continuum with two fiber families despite its known layered structure. This simplification is common in reproductive tract biomechanics: prior extension-inflation studies of the murine reproductive tract have generally used thin-walled pressure vessel theory to estimate average wall stress [25, 28], and human uterine iFEA studies have focused on a single dominant layer, the myometrium, given its primary load-bearing role [33]. While this simplification facilitates parameter identifiability, the reported constitutive parameters should be interpreted as effective homogenized properties that reproduce the experimentally observed deformation under the applied loading conditions.

The assumption of negligible axial strain was motivated by the large aspect ratio of the imaged uterine segment relative to its cross-sectional dimensions and the focus of the inverse analysis on cross-sectional deformation. Axial displacement measurements in a representative specimen supported this approximation, although it was not systematically evaluated across all specimens because quantifying axial deformation would have required a substantially larger imaging region of interest than that required to measure the in-plane deformations used for inverse parameter identification. Accordingly, the inverse analysis was based solely on cross-sectional boundary deformation, and the estimated material parameters were therefore informed primarily by radial and circumferential deformation. Consequently, the circumferential and longitudinal fiber families could not be independently identified from the available data. The two-family GOH formulation was retained to provide a complete representation of the known microstructural organization of the uterine wall, although the estimated fiber parameters are informed primarily by circumferential inflation kinematics. The estimated parameters also reflect quasi-static passive behavior and do not account for active smooth muscle contraction, viscoelasticity, or time-dependent tissue remodeling [11]. Finally, the fiber dispersion parameter was fixed throughout the optimization rather than estimated as a specimen-specific parameter, though its value was based on previous experimental results [9]. Future studies should evaluate the sensitivity of the inverse solutions to this modeling assumption.

Beyond the present validation study, this framework provides a foundation for investigating how biological remodeling processes alter uterine mechanics. Changes in collagen content, organization, crosslinking density, and extracellular matrix composition are known to occur during aging, disease progression, and exposure to environmental toxicants. Since the proposed approach links experimentally measured deformation to constitutive parameters with microstructural interpretations, it offers a means of quantifying the specific, tissue-scale mechanical consequences of such microstructural alterations. Future studies will employ the developed framework to examine reproductive tissue remodeling associated with environmental exposures, where changes in collagen organization and recruitment behavior are expected to influence organ-level mechanical function.

Despite these limitations, the proposed balloon inflation–based iFEA framework provides a robust and repeatable approach for estimating microstructure-informed constitutive parameters of soft hollow organs from experimentally measured deformation. The framework was sufficiently sensitive to detect mechanical changes induced by collagen crosslinking, demonstrating its potential for investigating how disease, aging, environmental exposures, or therapeutic interventions alter the mechanical behavior of reproductive tissues.

## Clinical trial number

Not applicable.

## Acknowledgements

This work was supported by NIH 5T32HD108075 and NIH 1R35ES034988. The research was carried out in the Beckman Institute for Advanced Science and Technology at the University of Illinois Urbana-Champaign. MicroCT imaging was performed using the Beckman Institute Microscopy Suite and computational analyses were performed using resources provided by the Beckman Institute Visualization Laboratory. Dr. Amy Wagoner Johnson, Dr. Indrani C. Bagchi, and Dr. Ayelet Ziv-Gal are Biohub Investigators.

## Data availability

The datasets generated and analyzed during the current study are available from the corresponding author upon reasonable request.

## Author contributions

Conceptualization: M.R. Arshee and A.J. Wagoner Johnson. Methodology: M.R. Arshee, C.M. Luetkemeyer, A.J. Wagoner Johnson, and A. Safar. Formal analysis: M.R. Arshee. Software: M.R. Arshee. Visualization: M.R. Arshee. Funding acquisition: J.A. Flaws, I.C. Bagchi, A. Ziv-Gal, and A.J. Wagoner Johnson. Supervision: C.M. Luetkemeyer, and A.J. Wagoner Johnson. Project administration: A.J. Wagoner Johnson. Writing—original draft: M.R. Arshee. Writing—review and editing: C.M. Luetkemeyer, J.A. Flaws, I.C. Bagchi, A. Ziv-Gal, and A.J. Wagoner Johnson. All authors read and approved the final manuscript.

