## Supplemental document for "Image-Informed Inverse Finite Element Analysis Reveals Altered Constitutive Behavior Following Controlled Uterine Tissue Remodeling"

Journal: Annals of Biomedical Engineering

**Mahmuda Raakib Arshee ^1^, Callan M. Luetkemeyer ^1,4,6,7,8^, Indrani C. Bagchi ^2,4,5^, Ayelet Ziv-Gal ^2,4,5^, Jodi A. Flaws ^2,4^, Adira Safar ^2^, A. J. Wagoner Johnson ^1,3,4,5,6,7,*^**

^1^ Mechanical Science and Engineering, Grainger College of Engineering, University of Illinois at Urbana-Champaign, Champaign, IL, 61820, USA

^2^ Comparative Biosciences, College of Veterinary Medicine, University of Illinois at Urbana-Champaign, Urbana, IL, 61802, USA

^3^ Biomedical and Translational Sciences, Carle Illinois College of Medicine, University of Illinois at Urbana-Champaign, Champaign, IL 61820, USA

^4^ Carl R. Woese Institute for Genomic Biology, University of Illinois at Urbana-Champaign, Urbana, IL, 61801, USA

^5^ Biohub Chicago, LLC, Chicago, IL, 60642, USA

^6^ Beckman Institute for Advanced Science and Technology, University of Illinois at Urbana-Champaign, Urbana, IL, 61801, USA

^7^ Department of Bioengineering, Grainger College of Engineering, University of Illinois at Urbana-Champaign, Urbana, IL, 61801, USA

^8^ Materials Research Laboratory, University of Illinois at Urbana-Champaign, Urbana, IL, 61801, USA

* Corresponding author at Mechanical Science and Engineering, Grainger College of Engineering, University of Illinois at Urbana-Champaign, Champaign, IL, 61820, USA

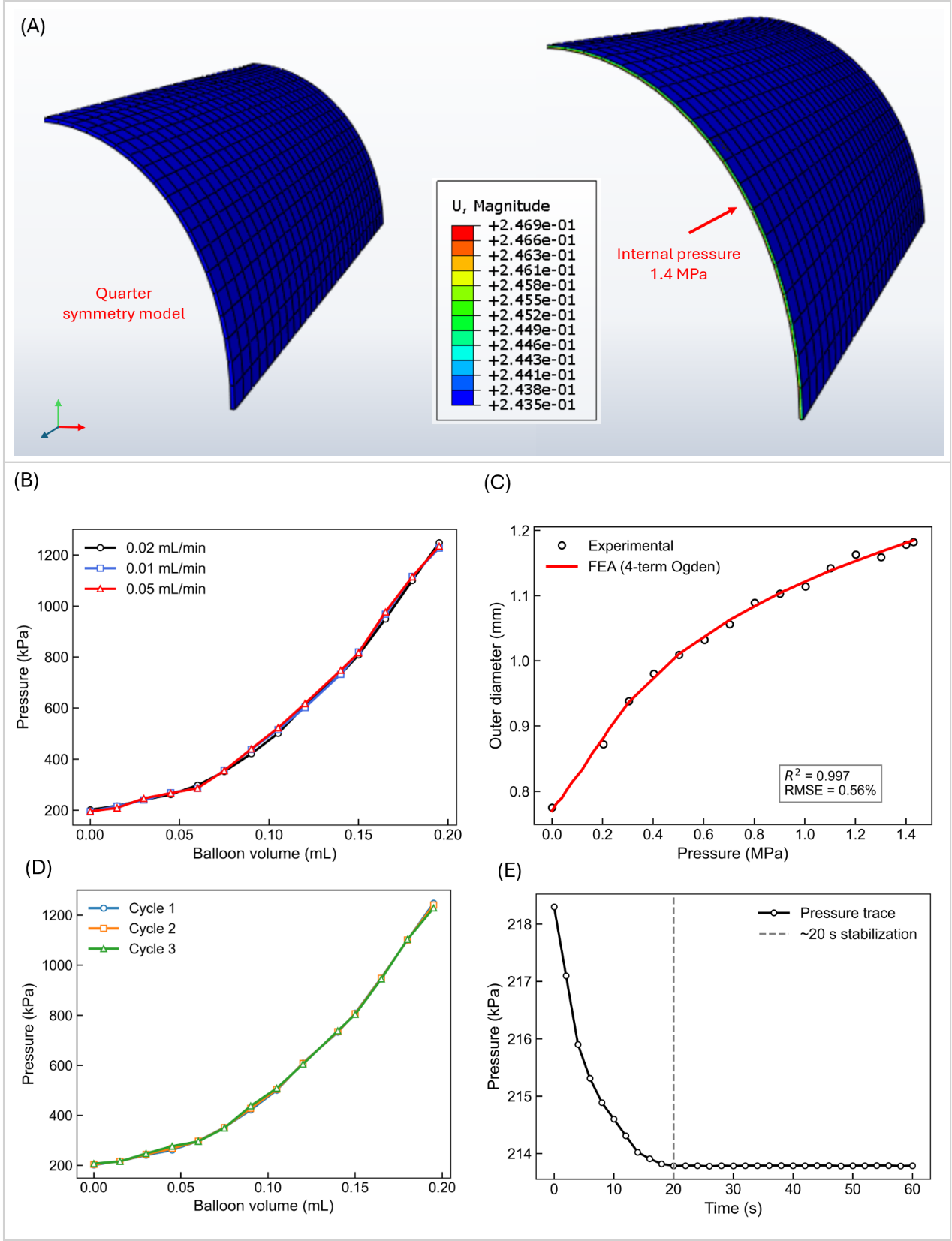

**Fig. S1** Balloon material model calibration and validation. Quarter-symmetry finite element model of the balloon in the undeformed (left) and inflated (right) configurations. The balloon was modeled using a four-term Ogden hyperelastic constitutive formulation and subjected to internal pressure. The displacement contour shown for the inflated configuration illustrates the deformation field at the maximum pressure level. (B) Pressure–volume responses of the balloon obtained at three inflation rates (0.01, 0.02, and 0.05 mL/min), demonstrating negligible dependence of balloon mechanics on the applied inflation rate over the range investigated. (C) Comparison between experimental and simulated balloon diameter–pressure responses following balloon material calibration by iFEA. (D) Pressure–volume responses from three consecutive balloon inflation cycles following preconditioning demonstrating repeatability of the balloon mechanical response. (E) Representative pressure stabilization following a single inflation increment. Measured tissue pressure reached equilibrium within approximately 20 s, supporting the use of a 1-min equilibration period before microCT image acquisition

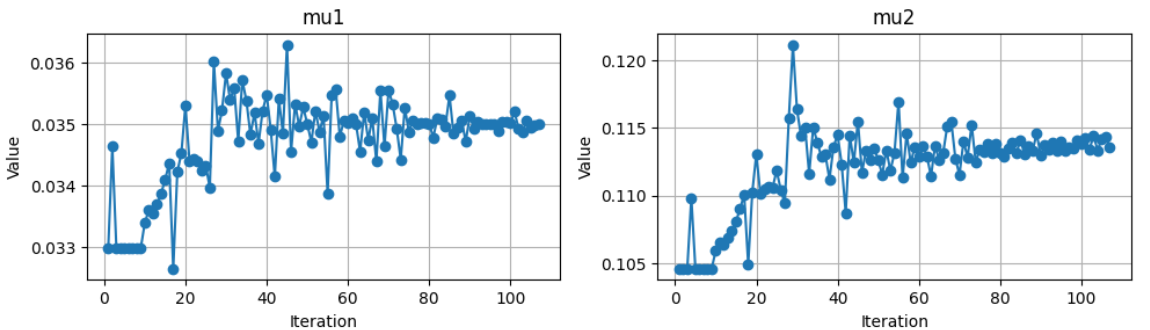

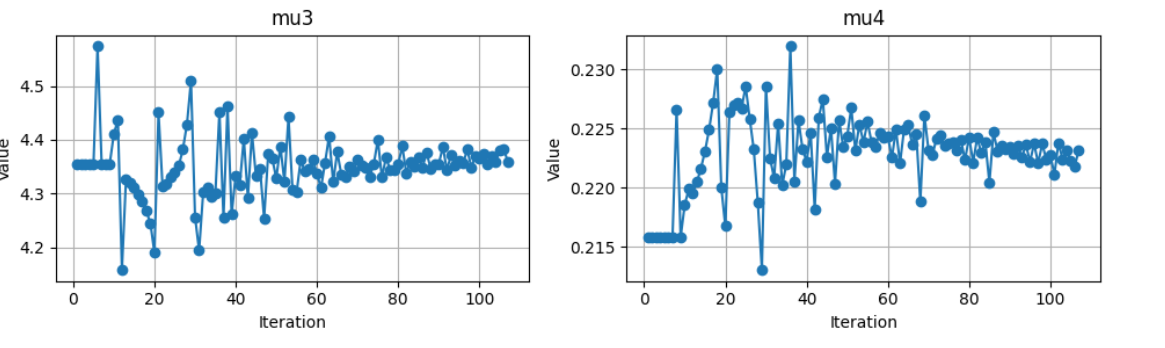

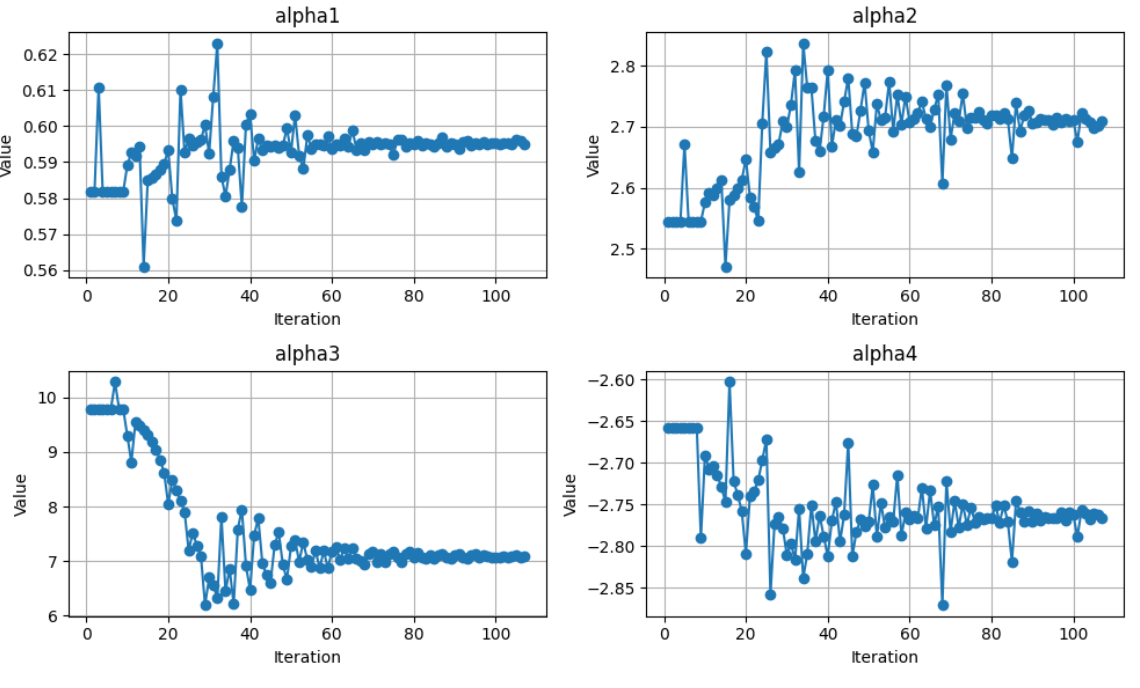

**Fig. S2** Optimization convergence of the balloon material parameters. Evolution of the optimized four-term Ogden material parameters during iFEA calibration of the balloon model. The parameters include four shear-modulus coefficients (μ₁–μ₄) and four strain-hardening exponents (α₁–α₄). All parameters converged toward stable values and the converged parameter set was used in all tissue inflation simulations

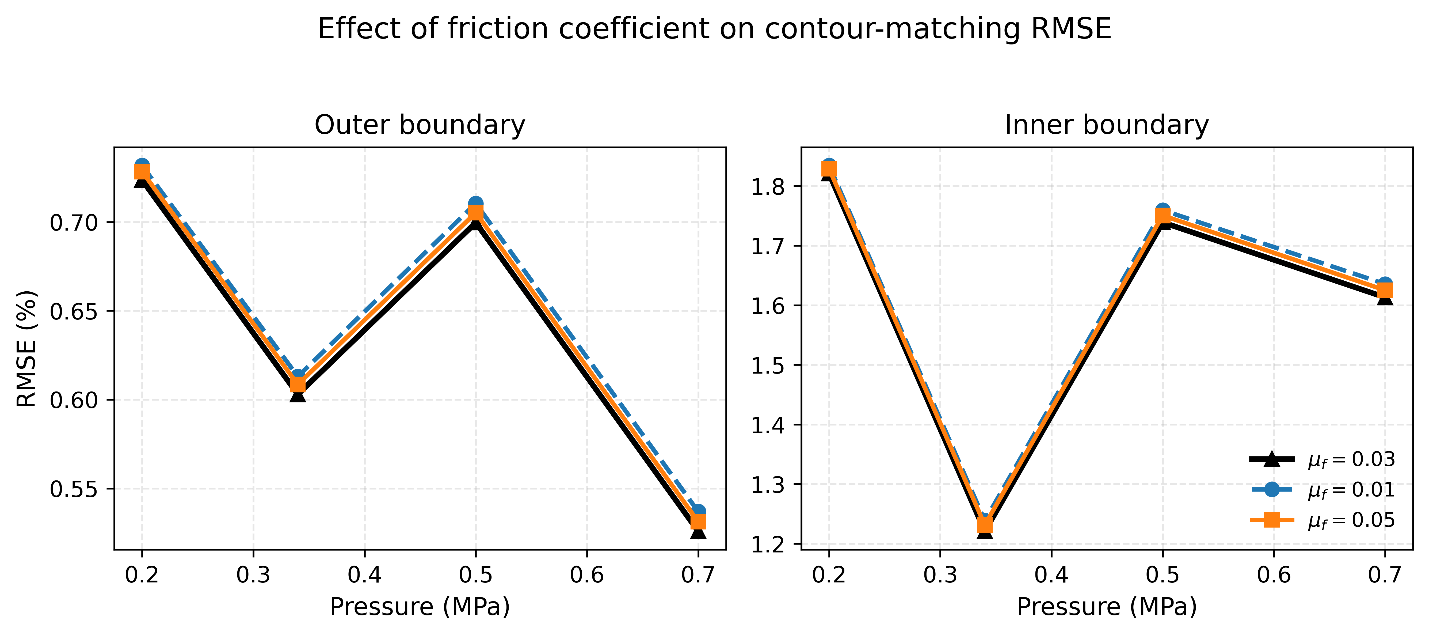

**Fig. S3** Effect of friction coefficient on contour-matching RMSE. Contour-matching RMSE obtained using friction coefficients of *μ_f_* = 0.01, 0.03, and 0.05 at multiple inflation pressures. (A) Outer boundary RMSE and (B) inner boundary RMSE. Changes in friction coefficient produced only minor variations in contour-matching RMSE across the investigated pressure range, with differences remaining much smaller than the overall fitting error. These results indicate that the iFEA framework is relatively insensitive to the assumed friction coefficient within the tested range, supporting the use of *μ_f_* = 0.03 in the final simulations

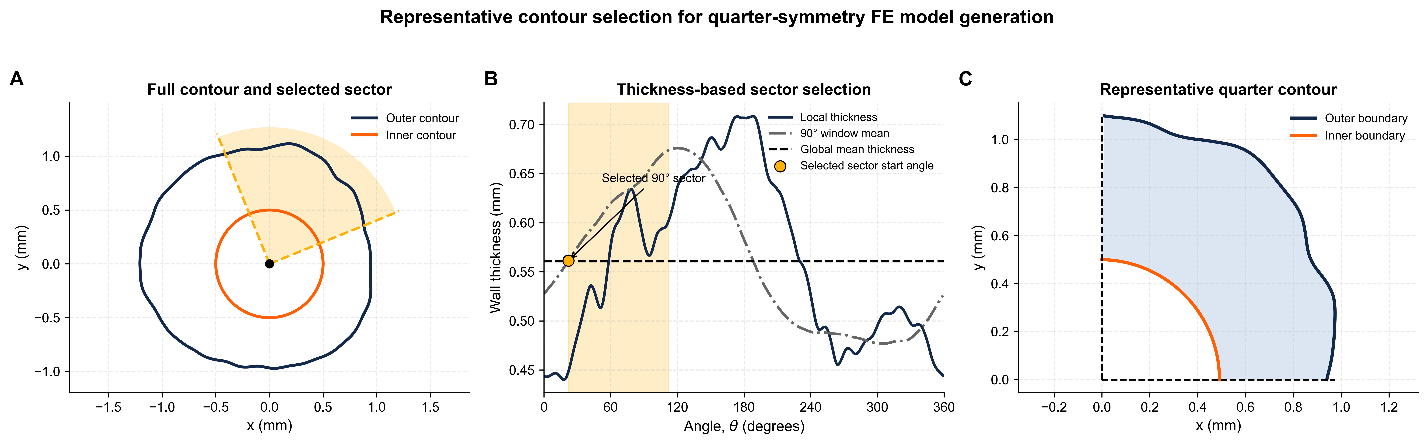

**Fig. S4** Selection of a representative tissue sector for finite element modeling. (A) Full cross-sectional contour of a representative uterine specimen showing the selected 90° sector used to construct the finite element model. The highlighted sector was chosen to represent the overall wall geometry to reduce computational costs. (B) Wall thickness as a function of circumferential position. The solid curve shows the local wall thickness around the specimen circumference, the dash-dotted curve shows the average thickness of candidate 90° sectors obtained using a sliding-window approach, and the horizontal dashed line indicates the global mean wall thickness of the full cross-section. The shaded region identifies the selected sector, whose average thickness most closely matches the global mean thickness. (C) Representative quarter contour generated from the selected sector and mapped into a first-quadrant geometry for finite element model construction

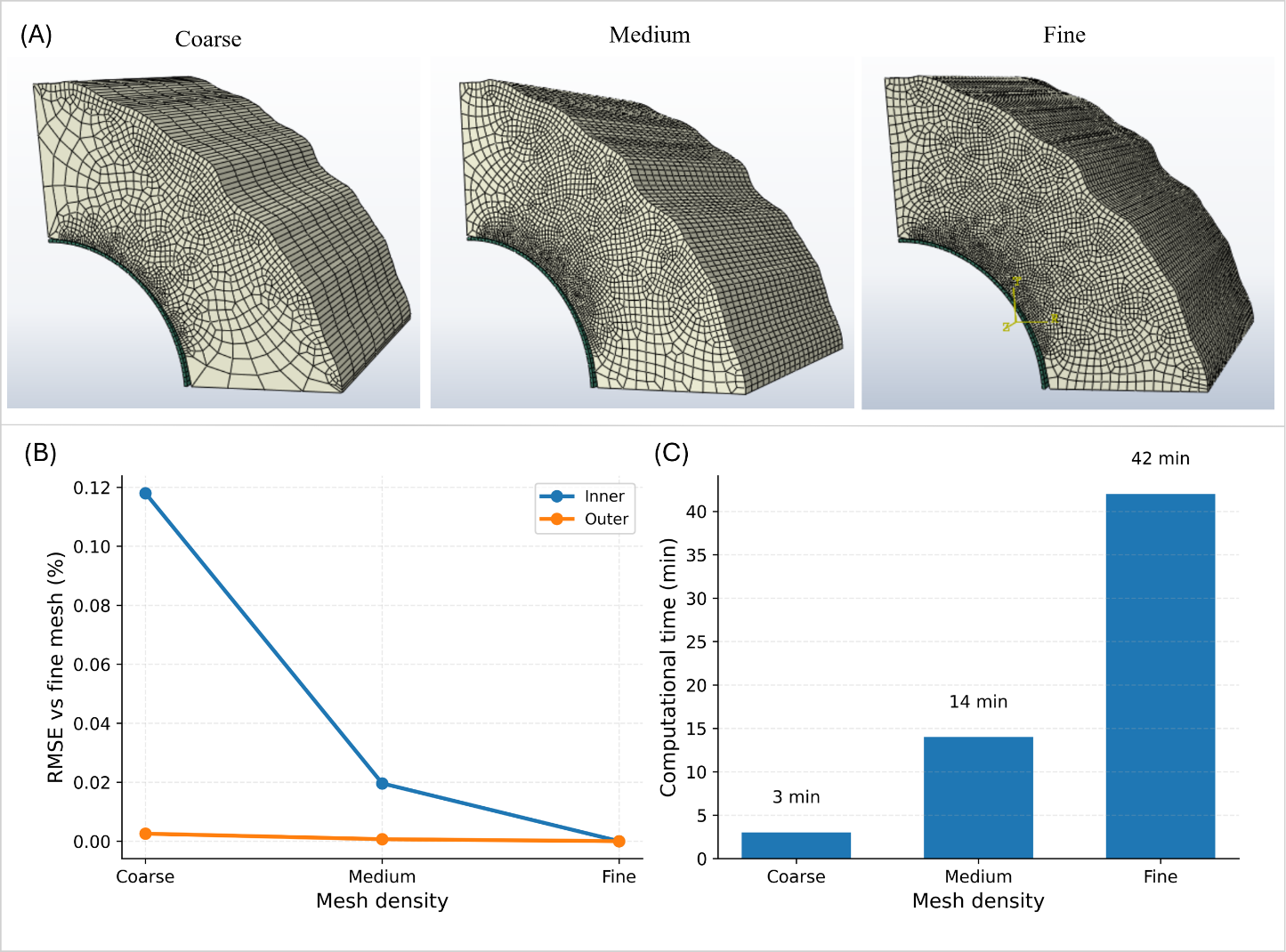

**Fig. S5** Mesh convergence and computational cost analysis. (A) Coarse, medium, and fine finite element meshes used for convergence assessment. (B) RMSE of the coarse and medium mesh solutions relative to the fine-mesh solution, calculated separately for the inner and outer contours. The medium mesh produced results nearly identical to the fine mesh, while the coarse mesh exhibited larger deviations. (C) Computational time associated with each mesh density. Based on the tradeoff between accuracy and computational cost, the medium mesh was selected for all subsequent simulations

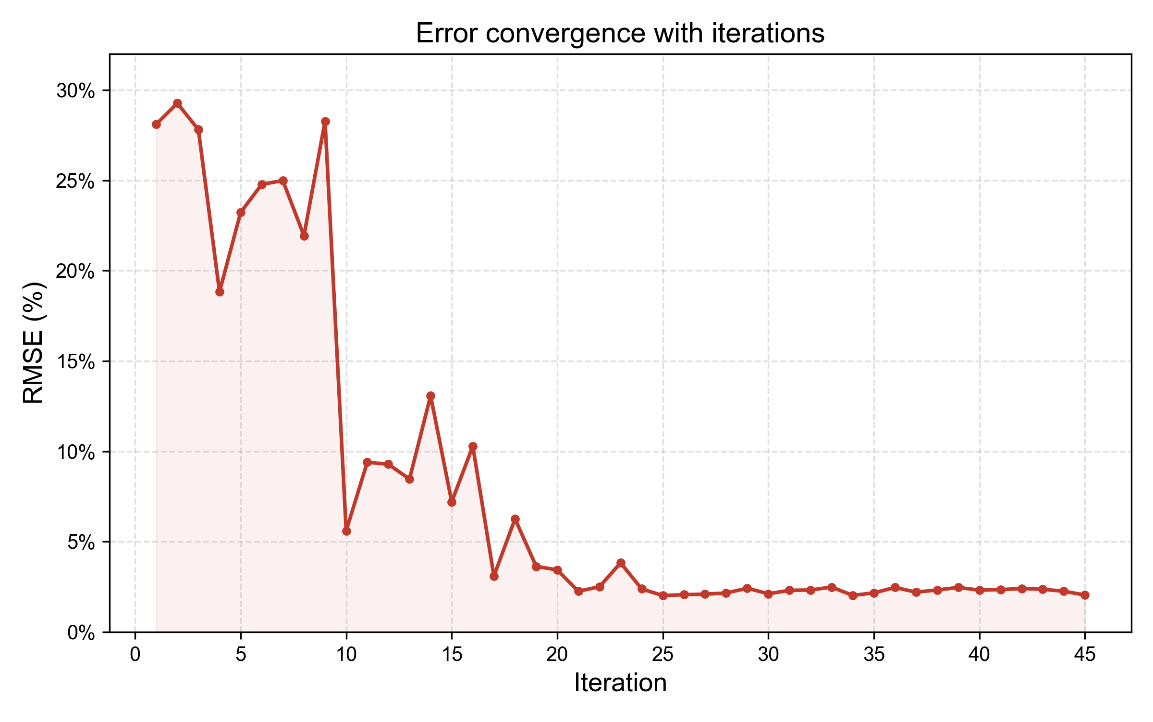

**Fig. S6** Representative optimization convergence during iFEA. Evolution of the contour-matching RMSE during optimization for a representative sample. RMSE between simulated and experimental contours is shown as a function of iteration number. The optimization rapidly reduced the fitting error during the early iterations, followed by convergence toward a stable minimum. Final RMSE values stabilized near 2%, indicating convergence of the iFEA parameter identification procedure

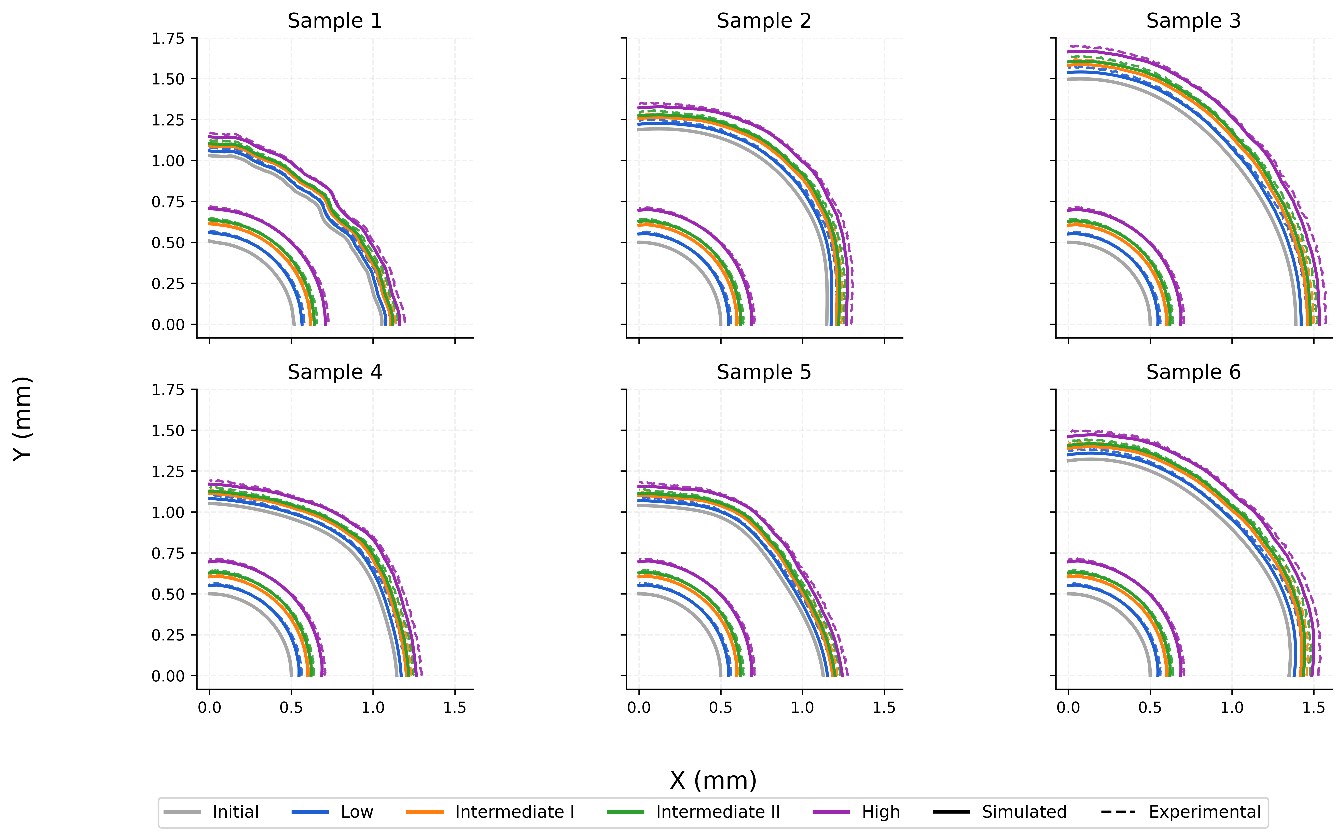

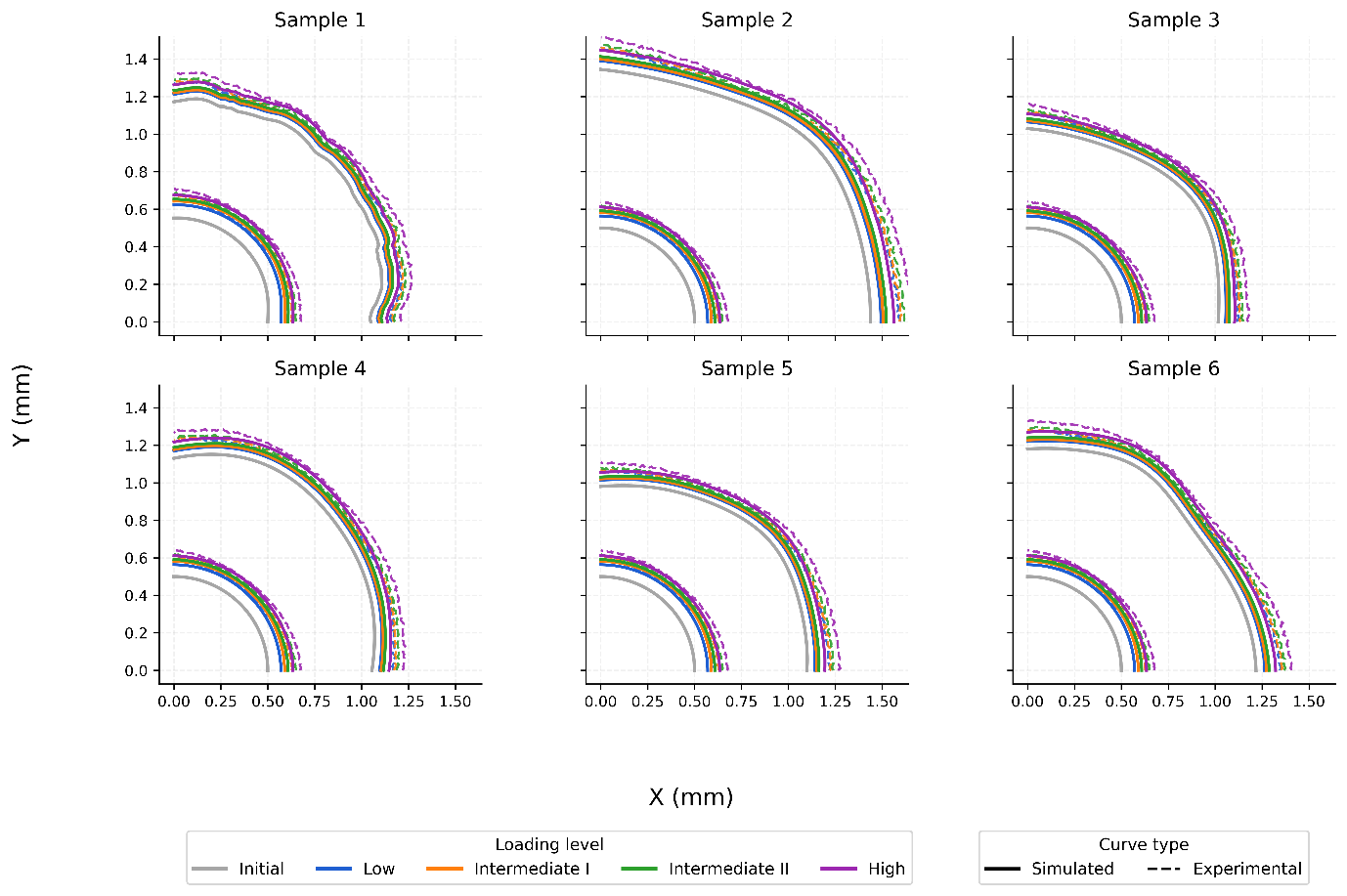

**Fig. S7** Contour-matching validation results for all uterine samples. Comparison of simulated and experimental tissue contours at multiple inflation pressures for all untreated (top) and GA-treated (bottom) samples (n = 6 per group). Solid lines represent finite element predictions and dashed lines represent experimentally measured contours. Gray curves indicate the initial undeformed geometry, while colored contours correspond to increasing inflation pressures (low, intermediate I, intermediate II, and high)
